# Cyclic electron transport via the NDH complex sustains photosynthesis and productivity under fluctuating and sub-optimal environments

**DOI:** 10.64898/2026.04.02.716017

**Authors:** Hiromasa Kodama, Wataru Yamori

## Abstract

The chloroplast NADH dehydrogenase-like (NDH) complex mediates cyclic electron transport (CET) around photosystem I (PSI) and contributes to photosynthetic regulation and photoprotection under various environmental stresses. Although NDH function has been extensively characterized under controlled conditions, NDH-deficient mutants often show only subtle phenotypes in such environments, leaving its physiological importance under naturally fluctuating field conditions poorly understood. Here, we evaluated growth, yield, and photosynthetic performance of NDH-deficient rice cultivated under field conditions. Mutant plants exhibited reduced biomass accumulation and grain yield compared with wild type. Detailed physiological analyses revealed that NDH deficiency markedly decreased PSI electron transport and CO₂ assimilation, particularly under low temperature and sub-saturating irradiance. At moderate and high temperatures, reductions in carbon fixation were largely confined to low-light conditions, whereas at low temperatures, impairment extended across nearly the entire light response range. Under repetitive fluctuating light regimes, NDH-deficient plants showed progressive declines in photosynthesis accompanied by a selective decrease in PSI photochemical capacity without changes in PSII maximum efficiency, indicating PSI-specific photoinhibition. These findings demonstrate that NDH-dependent CET plays a crucial role in sustaining photosynthetic efficiency and crop productivity in dynamic field environments by stabilizing PSI redox balance and maintaining long-term carbon gain.

**Summary Statement:** NDH-dependent cyclic electron transport supports photosynthesis and yield in field-grown rice by maintaining PSI function under fluctuating light, low temperature, and sub-saturating irradiance.

## Introduction

In natural environments, plants are continuously exposed to dynamic fluctuations in light intensity, spectral quality, and temperature that occur across timescales ranging from seconds to seasons. Such environmental variability strongly influences photosynthetic performance and ultimately determines plant productivity and crop yield (Pearcy, 1990; Long et al., 2006; Yamori, 2016). In particular, rapid increases in irradiance caused by sunflecks or canopy gaps can induce transient over-reduction of the photosynthetic electron transport chain, especially on the acceptor side of photosystem I (PSI), resulting in the generation of reactive oxygen species (ROS) and PSI-specific photoinhibition (Yamori et al., 2016a; Kono et al., 2017; Miyake, 2020). Conversely, periods of low light or suboptimal temperature can limit ATP supply and carbon assimilation capacity, leading to imbalances between energy absorption and utilization (Yamori et al., 2011, 2016a). Therefore, plants must maintain a delicate balance between maximizing photosynthetic efficiency and preventing photodamage under naturally fluctuating environmental conditions (Tikkanen & Grebe, 2018).

A central regulatory mechanism enabling such acclimation is cyclic electron transport (CET) around PSI. CET redirects electrons from ferredoxin (Fd) back to the plastoquinone (PQ) pool via the cytochrome *b*_6_/*f* complex, thereby generating proton motive force (*pmf*) and promoting ATP synthesis without net NADPH production (Yamori & Shikanai, 2016; Shikanai et al., 2025). This process contributes to multiple aspects of photosynthetic regulation, including lumen acidification, activation of non-photochemical quenching (NPQ), and control of electron flux toward PSI (Yamori & Shikanai, 2016; Shikanai & Yamamoto, 2017). Through these mechanisms, CET plays a critical role in sustaining photosynthesis and plant growth under various environmental constraints.

In angiosperms, CET is mediated by at least two distinct pathways: one dependent on the PROTON GRADIENT REGULATION5 (PGR5)/PGR5-LIKE1 (PGRL1) protein complex and another involving the chloroplast NADH dehydrogenase-like (NDH) complex (Shikanai et al., 1998; Munekage et al., 2004; DalCorso et al., 2008). The PGR5-dependent pathway has been extensively studied and is known to be essential for photoprotection under high light and fluctuating light conditions, as evidenced by the severe growth defects of *pgr5* mutants in *Arabidopsis thaliana* exposed to dynamic irradiance regimes (Tikkanen et al., 2010; Suorsa et al., 2012). In contrast, the physiological significance of NDH-dependent CET has long remained less clear, partly because NDH-deficient mutants often show only subtle phenotypes under steady laboratory conditions (Peng & Shikanai, 2011; Wang et al., 2015).

Recent studies have begun to clarify that the NDH complex contributes to photosynthetic regulation in specific environmental contexts. The NDH complex accepts electrons from reduced ferredoxin and transfers them to PQ while simultaneously translocating protons across the thylakoid membrane, thereby enhancing the formation of ΔpH and modulating *pmf* partitioning (Yamori et al., 2015, 2016a; Basso et al., 2020). This proton-pumping activity supports photosynthetic control by regulating electron transport at the cytochrome *b*_6_*f* complex, particularly during photosynthetic induction and under low-light conditions (Yamori et al., 2015; Basso et al., 2020; Zhou et al., 2023). Moreover, NDH has been shown to functionally interact with ion transporters such as KEA3, highlighting its role in fine-tuning the balance between photoprotection and photosynthetic efficiency (Armbruster et al., 2014; Basso et al., 2020).

Importantly, NDH-dependent CET is especially relevant in angiosperms that lack flavodiiron proteins (FLVs), which provide alternative electron sinks in cyanobacteria, mosses, and some gymnosperms (Rantala et al., 2020). In major crop species such as rice and Arabidopsis, CET pathways therefore represent the primary mechanisms preventing PSI over-reduction and photoinhibition. Consistent with this notion, NDH deficiency has been shown to reduce PSI electron transport capacity and CO₂ assimilation under low temperature, low light, and fluctuating light in controlled growth chamber experiments (Yamori et al., 2011, 2015, 2016a). NDH-dependent CET has also been implicated in plant responses to diverse abiotic stresses, including drought, heat, and high irradiance (Horváth et al., 2000; Rumeau et al., 2007; Zhang & Sharkey, 2009).

However, most previous investigations of CET function have relied on simplified and artificial environmental regimes, such as repetitive stepwise transitions between high and low light. Increasing evidence suggests that such experimental designs may not adequately capture the complexity of natural field environments (Lawson et al., 2012; Vialet-Chabrand et al., 2017). Indeed, plants grown under light regimes that mimic natural fluctuations exhibit distinct photosynthetic acclimation patterns compared with those grown under constant or monotonous fluctuating light conditions (Burgess et al., 2023). Despite the growing recognition that realistic environmental variability must be considered to understand photosynthetic regulation in crops, the functional significance of NDH-dependent CET under actual field conditions remains largely unexplored.

To address this knowledge gap, we investigated the physiological role of the NDH complex in rice (*Oryza sativa* L.) cultivated under natural field environments. By combining growth analysis, yield evaluation, and detailed measurements of photosynthetic responses across a range of temperatures and light intensities, we aimed to determine how NDH-dependent CET contributes to maintaining PSI function and carbon assimilation under realistic environmental stresses. Our results demonstrate that NDH plays a critical role in sustaining photosynthetic efficiency, preventing PSI photoinhibition, and supporting crop productivity under naturally variable conditions. These findings provide direct field-based evidence that NDH-dependent CET is an essential regulatory mechanism enabling crops to cope with the complex and dynamic environments encountered in agricultural systems.

## Materials & methods

### Plant materials and growth conditions

The rice mutant defective in the OsCRR6 gene (Os08g0167500) due to the insertion of the Tos17 retrotransposon (*crr6; −/−*), its wild type (*Oryza sativa* ssp. *japonica* cv. Hitomebore), and control progeny lacking the Tos17 insertion in *OsCRR6* but sharing the same genetic background as the homozygous mutant lines (control*; +/+*) were used in this study (Yamori et al. 2015, 2016a).

Plants were grown under field conditions at the Institute for Sustainable Agro-ecosystem Services, Graduate School of Agricultural and Life Sciences, The University of Tokyo (Tokyo, Japan) from late April to early October 2020. Environmental conditions during the experimental period are shown in a Figure.

Plants were sampled at 60 and 160 days after germination. Leaf shoots and roots were oven-dried at 80°C, and their dry weights were determined. At 160 days, plants were harvested and air-dried prior to sampling. Grain weight was subsequently measured to determine yield.

### Photosynthesis analysis

Gas exchange, chlorophyll fluorescence, and P700 redox state were measured simultaneously using a GFS-3000 gas-exchange system and a Dual-PAM-100 measuring system (Walz, Germany) in the uppermost fully expanded leaves of 60–80-day-old plants (Yamori et al 2016a; Qu et al. 2025). After 30 min dark adaptation, a saturating pulse was applied to determine maximum fluorescence and the maximal P700 signal. Photosynthetic parameters were recorded every 20 s at a CO₂ concentration of 400 µmol mol⁻¹ under either constant high light or fluctuating light conditions. The quantum yield of PSI (Y(I)) was calculated as Y(I) = 1 – Y(ND) – Y(NA), where Y(ND) corresponds to the fraction of P700 that is already oxidized by actinic light and Y(NA) corresponds to the fraction of P700 that is closed owing to acceptor side limitation (Klughammer & Schreiber, 1994). The quantum yield of photosystem II [Y(II) = (*F*_m_′–*F*′)/*F*_m_′], photochemical quenching [qP = (*F*_m_′–*F*′)/(*F*_m_′–*F*_o_′)], non-photochemical quenching [NPQ = (*F*_m_–*F*_m_′)/*F*_m_′] and the fraction of PSII centers in the open state (with plastoquinone oxidized) [qL = qP × (*F*_o_′/*F*′)] were calculated (Genty et al., 1989; Hendrickson et al., 2004; Kramer et al., 2004). The electron transport rate (ETR) was calculated as ETR I (or ETR II) = 0.5 × abs I × Y(I) (or Y(II)), where 0.5 is the fraction of absorbed light reaching PSI or PSII and abs I is absorbed irradiance taken as 0.84 of incident irradiance.

### Determination of leaf biochemical and protein contents

Immediately after gas-exchange measurements, leaf samples were collected, frozen in liquid nitrogen, and stored at −80°C. Frozen samples were ground in liquid nitrogen and homogenized in extraction buffer. Leaf nitrogen, chlorophyll, and Rubisco contents were quantified as described by Yamori et al. (2011). Proteins were separated by SDS-PAGE and transferred to polyvinylidene difluoride membranes. The abundance of CRR6, cytochrome f (cytochrome *b_6_*/*f* complex), and NdhK (a subunit of NDH subcomplex A) was determined using specific antibodies. A dilution series of wild-type protein extracts was included to estimate relative protein abundance in mutant samples.

Rubisco large subunit content was determined spectrophotometrically following formamide extraction of Coomassie Brilliant Blue R-250-stained bands corresponding to Rubisco subunits. Chlorophyll was extracted in 80% (v/v) acetone and quantified according to Porra et al. (1989). Leaf nitrogen content was measured using a CHNO/S elemental analyzer (Vario EL III, Elementar, Germany).

### Blue-Native PAGE (BN-PAGE) analysis

BN-PAGE was performed as described by Peng et al. (2010) with minor modifications. Freshly isolated thylakoid membranes were gently washed twice with buffer containing 25 mM BisTris-HCl (pH 7.0) and 20% glycerol, and then solubilized in the same buffer supplemented with 1.25% (w/v) n-dodecyl-β-D-maltoside (DDM) at a final chlorophyll concentration of 1 mg mL⁻¹. After incubation on ice for 10 min, samples were centrifuged at 12,000 × g for 10 min. The supernatants were supplemented with one-tenth volume of BN sample buffer (100 mM BisTris-HCl pH 7.0, 5% Serva Blue G, 0.5 M 6-amino-n-caproic acid, and 30% sucrose). Equal amounts of chlorophyll were loaded per lane. Following electrophoresis, thylakoid protein complexes were visualized by staining with Coomassie Brilliant Blue (CBB).

### Analysis of photoinhibition

Leaves were placed in a temperature-controlled chamber at 400 µmol mol⁻¹ CO₂ and 65% relative humidity within the Dual-PAM-100 and GFS-3000 systems. Leaves were exposed to fluctuating light consisting of alternating high light (1500 µmol m⁻² s⁻¹) and low light (200 µmol m⁻² s⁻¹) at cycle lengths of 10, 5, or 2 min for 5 h (10-min HL/LL cycles , 5-min HL/LL cycles , 2-min HL/LL cycles). The maximal P700 signal (Pm) and maximum PSII quantum yield (Fv/Fm) after 30 min dark incubation were measured before and after treatment. Photoinhibition was evaluated as the reduction in Pm and Fv/Fm following fluctuating light exposure.

### Statical analyses

Statistical analyses were conducted using R software (version 4.1.2). One-way ANOVA was performed to compare differences between WT and transgenic plants, followed by Tukey’s post-hoc test to identify significant pairwise differences. Statistical significance was set at *p* < 0.05 for all analyses.

## Results

### Loss of NDH activity in the *crr6* mutant

The chloroplast NDH complex consists of five subcomplexes (SubA, SubB, SubE, SubM, and SubL). CRR6 is required for the assembly of NdhI in subcomplex A (Shikanai et al., 2025). Immunoblot analysis confirmed that CRR6 protein was absent in the *crr6* mutant (Fig. 1A). To examine the effect of CRR6 deficiency on NDH accumulation, we analyzed the level of NdhK, a subunit of subcomplex A. NdhK was not detected in the *crr6* mutant, consistent with previous observations (Yamori et al., 2015).

**Figure 1.**
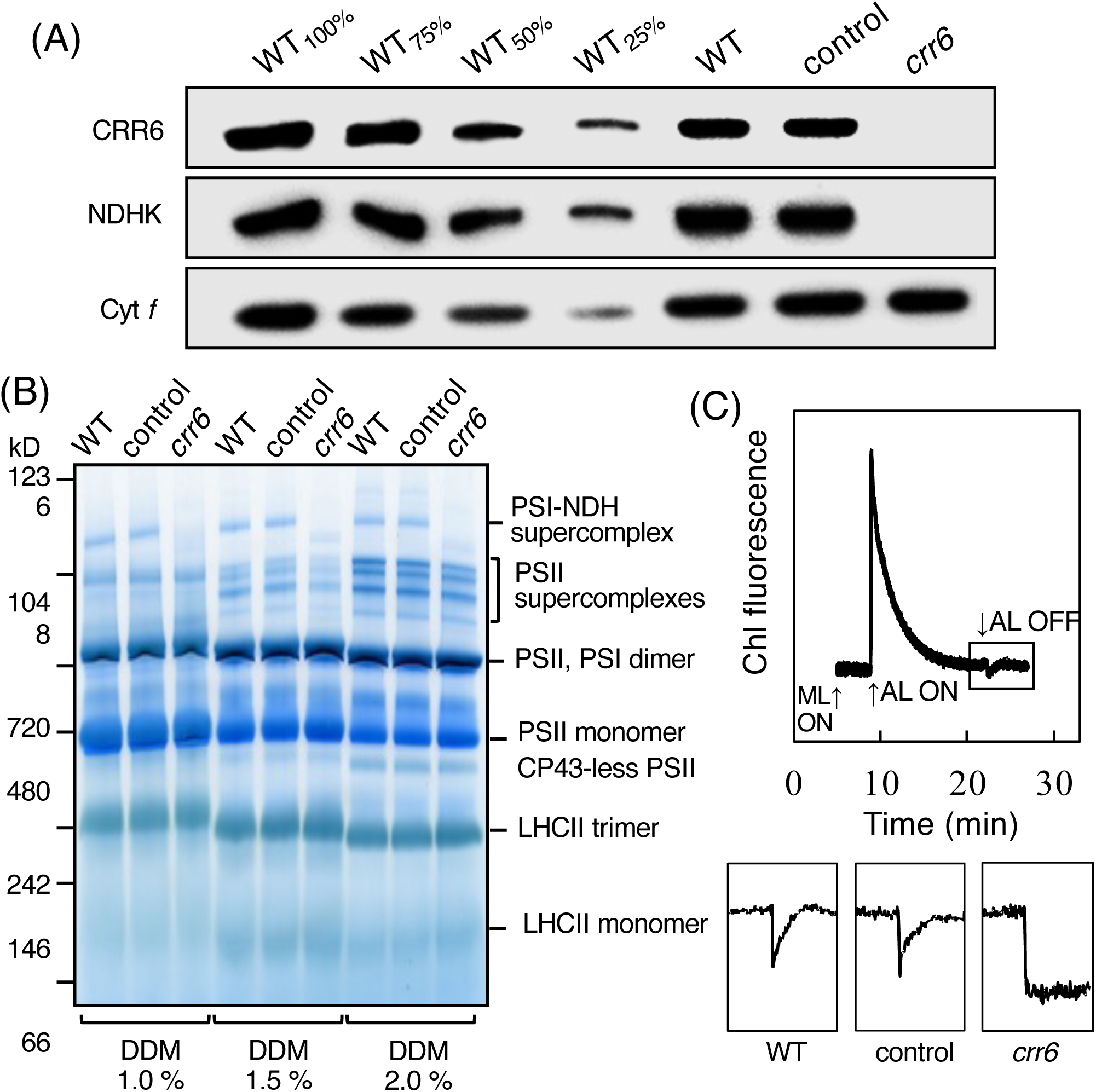
Characterization of the NDH-deficient *crr6* plants. (A) Thylakoid membrane protein complexes isolated from wild-type (WT), control, and crr6 plants. Thylakoid membranes were solubilized with 1.0%, 1.5%, or 2.0% (w/v) dodecyl maltoside and separated by blue native-PAGE. Gels were stained with Coomassie Brilliant Blue. Equal amounts of chlorophyll were loaded per lane. (B) NDH activity monitored by chlorophyll fluorescence analysis. A representative fluorescence trace from WT plants is shown. Arrows indicate the timing of measuring light (ML) and actinic light (AL; 200 μmol m⁻² s⁻¹). The transient increase in fluorescence following AL cessation (boxed region) was used as an indicator of NDH activity. Enlarged traces from WT, control, and *crr6* plants are shown below. (C) Immunoblot analysis of photosynthetic proteins. Total leaf proteins were separated by SDS-PAGE and detected using the indicated antibodies. Proteins were loaded on an equal fresh-weight basis. CRR6 is a stromal protein required for accumulation of NDH subcomplex A, whereas NDHK is a subunit of this subcomplex. Cytochrome f (Cyt *f*) is a component of the cytochrome b₆f complex.

In angiosperms, the NDH complex forms a supercomplex with two PSI complexes, which can be detected as a high-molecular-weight band by BN-PAGE. This PSI–NDH supercomplex was observed in WT and control plants but was absent in the *crr6* mutant (Fig. 1B).

NDH activity was further assessed by monitoring the transient post-illumination increase in chlorophyll fluorescence, which reflects NDH-dependent reduction of the plastoquinone pool in darkness (Shikanai et al., 1998). This fluorescence rise was detected in WT and control plants but was completely absent in the *crr6* mutant (Fig. 1C), demonstrating a complete loss of NDH activity.

### NDH-dependent CET contributes to plant growth and yield under field conditions

Plants were cultivated in the field from April 26 to October 4 under naturally fluctuating light and temperature conditions (Fig. 2A, B). The *crr6* mutant showed significantly reduced dry mass at both 60 and 160 days after germination compared with WT and control plants (Fig. 2C, D). Grain yield was also significantly lower in the crr6 mutant (Fig. 2E).

**Figure 2.**
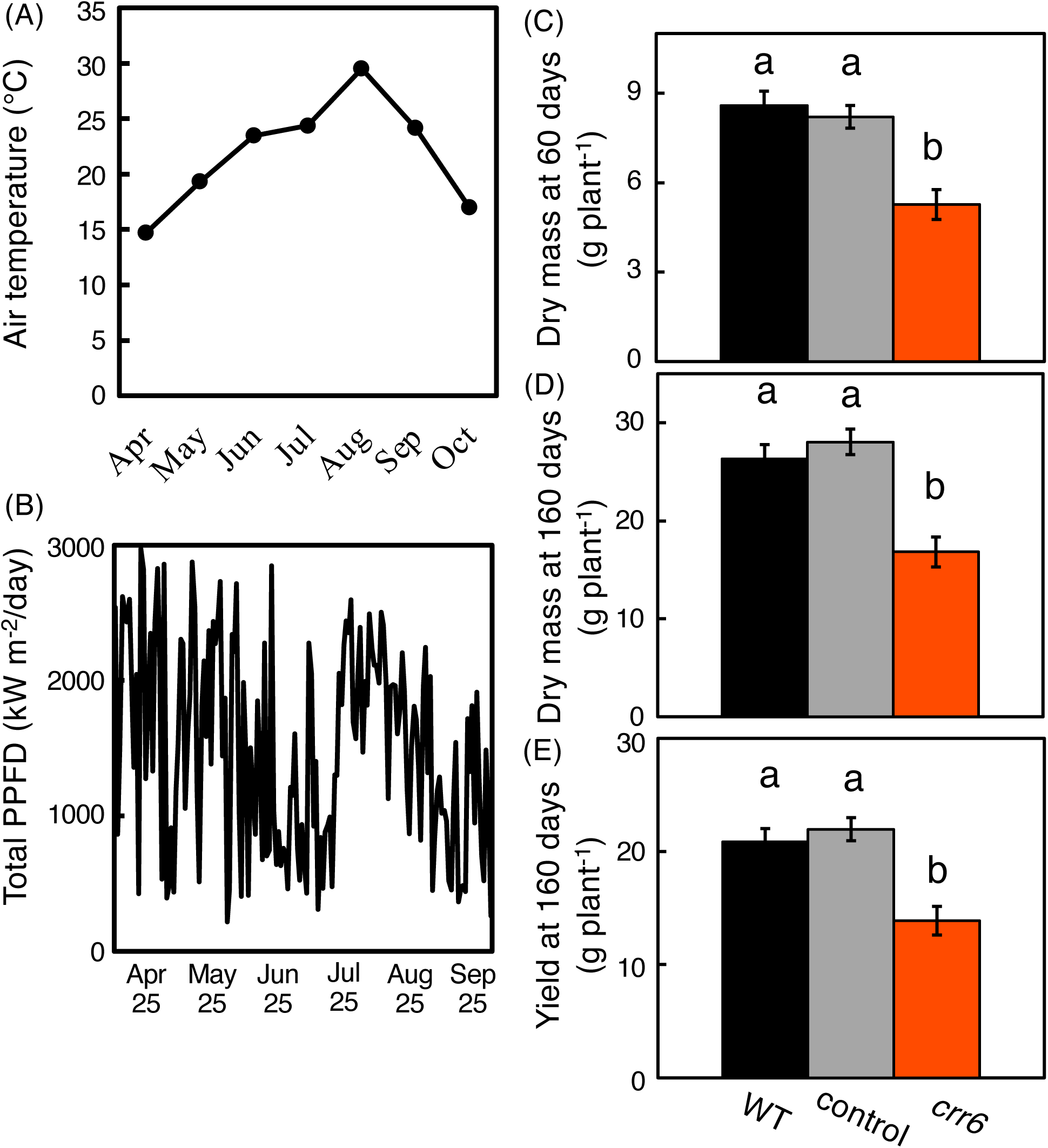
Growth and yield responses of *crr6* plants under outdoor cultivation. Seedlings were cultivated from 26 April to 4 October in a greenhouse under natural environmental conditions. (A, B) Monthly mean air temperature and daily cumulative photosynthetic photon flux density (PPFD) during the cultivation period. (C, D) Shoot and root dry mass measured at 60 days (C) and 160 days (D) after transplanting. (E) Grain yield at 160 days. Values represent means ± SE (n = 3–5). Different letters indicate significant differences among WT, control, and crr6 plants according to Tukey–Kramer multiple comparison tests (*p* < 0.05).

### Leaf biochemical properties were unaffected by NDH deficiency

Leaf nitrogen content, Rubisco content, cytochrome f abundance, and chlorophyll content per unit leaf area were similar among WT, control plants, and the *crr6* mutant (Table 1). The chlorophyll *a*/*b* ratio was also comparable between genotypes (Table 1), indicating that NDH deficiency did not alter major biochemical leaf traits.

**Table 1.**
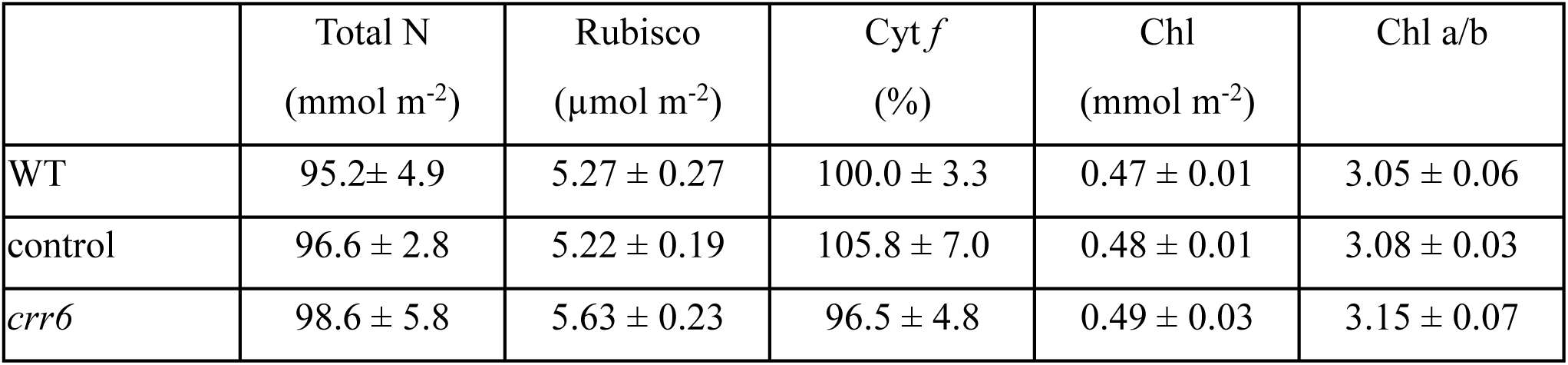
Effect of the *crr6* defect on physiological components of photosynthesis. Contents of total nitrogen (Total N), Rubisco, Cytochrome *f* (Cyt *f*), Chlorophyll (Chl) and the ratio of Chlorophyll *a* and *b* (Chl *a*/*b*) were quantified. The Cyt *f* content is shown in a percentage relative to WT plants. Data represent means ± SE, n = 3∼5. As a result of Tukey–Kramer multiple comparison tests (*p* < 0.05), there was no difference between WT, control plants, and *crr6* mutant.

### NDH-dependent CET supports PSI activity and CO₂ assimilation under low light and low temperature

Light-response curves of photosynthetic parameters were measured at 18, 28, and 38°C (Figs. 3–5). NDH deficiency had no significant effect on stomatal conductance (*g_s_*), intercellular CO₂ concentration (*C_i_*), or dark respiration rate (*Rd*) across temperatures and light intensities (Fig. 4D–I; Fig. 5G). However, the *crr6* mutant exhibited significantly lower CO₂ assimilation rates under relatively weak light, although the affected light range depended on temperature. At 28 °C and 38 °C, differences were observed between 200 and 1000 µmol m⁻² s⁻¹, whereas at 18 °C, reductions occurred across nearly the entire light range (Fig. 4A–C). ETRII was largely similar between genotypes except at moderate light intensities (500–750 µmol m⁻² s⁻¹) at 18°C (Fig. 3J–L; Fig. 5E, F). In contrast, ETRI was significantly lower in the *crr6* mutant under 1000 µmol m⁻² s⁻¹ across all temperatures (Fig. 3A–C; Fig. 5C, D).

**Figure 3.**
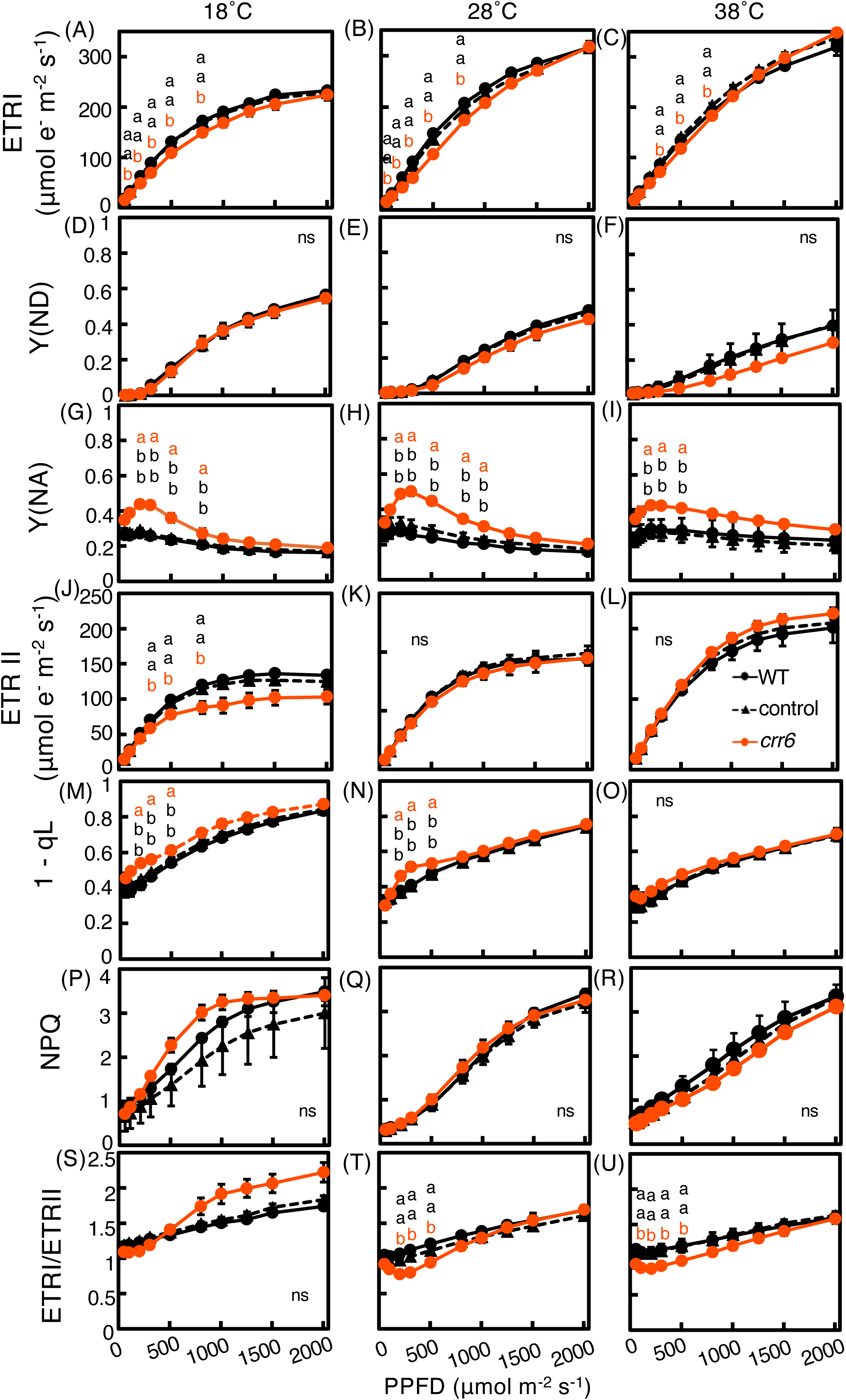
Effects of NDH deficiency on PSI redox state and PSII fluorescence parameters across light intensities at different temperatures. Light-response curves of PSI redox parameters were determined at 18, 28, and 38 °C. (A–C) Electron transport rate through PSI [ETRI]. (D–F) Donor-side limitation of PSI [Y(ND)]. (G–I) Acceptor-side limitation of PSI [Y(NA)]. (J–L) Electron transport rate through PSII [ETRII]. (M–O) Fraction of closed PSII centers (1 − qL). (P–R) Non-photochemical quenching (NPQ). (S–U) Ratio of ETRI to ETRII. Values represent means ± SE (n = 3–5). Significant differences among genotypes were evaluated using Tukey–Kramer tests (*p* < 0.05). Different letters indicate significant differences among WT, control, and *crr6* plants according to Tukey–Kramer multiple comparison tests (*p* < 0.05). black symbols represent WT and control plants, while orange symbols represent *crr6* mutant.

**Figure 4.**
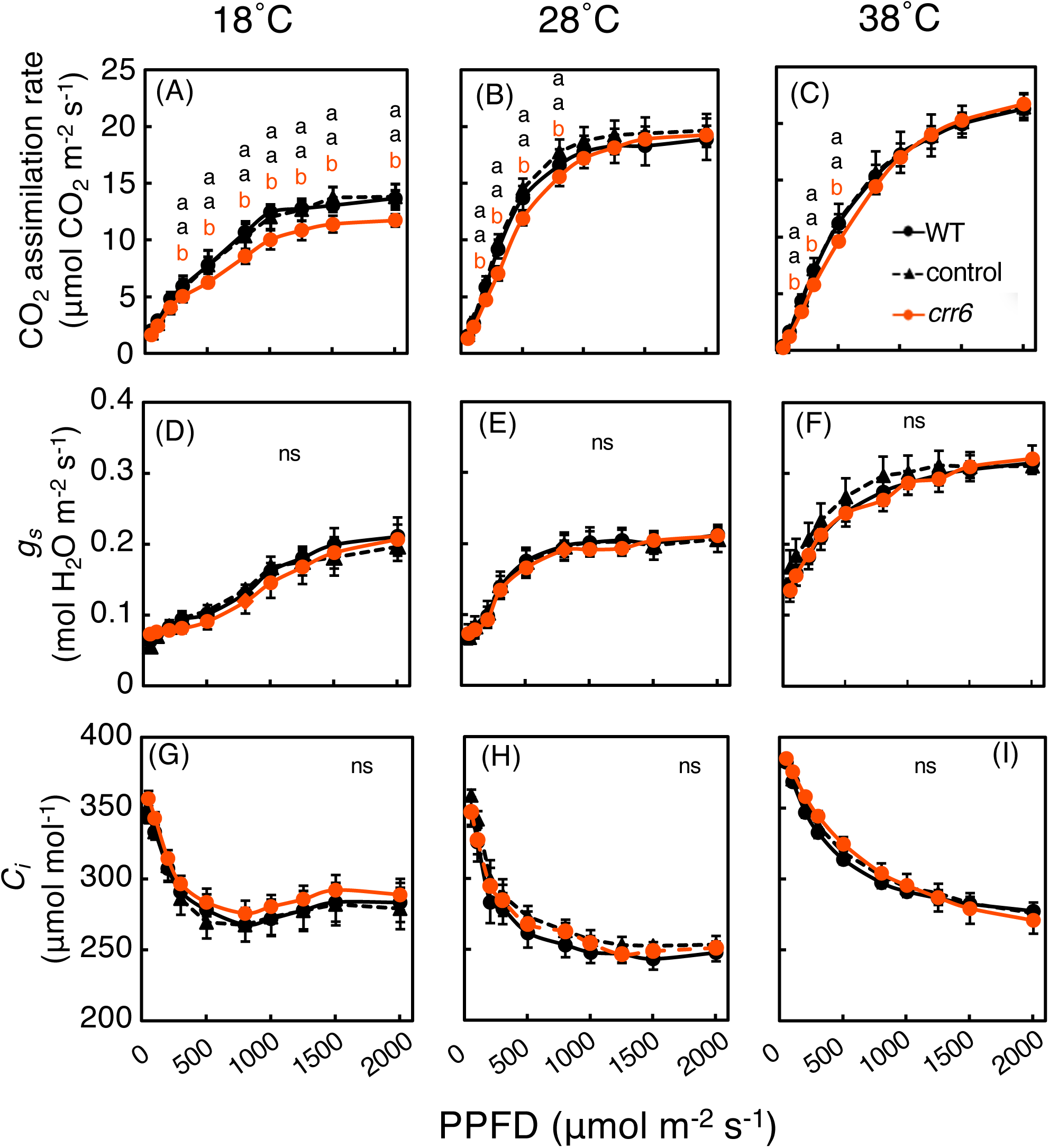
Effects of NDH deficiency on gas-exchange responses across light intensities at different temperatures. Light-response curves of gas-exchange parameters were determined at 18, 28, and 38 °C. (A–C) Net CO₂ assimilation rate. (D–F) Stomatal conductance (gₛ). (G–I) Intercellular CO₂ concentration (Cᵢ). Values represent means ± SE (n = 3–5). Significant differences among genotypes were evaluated using Tukey–Kramer tests (*p* < 0.05). Different letters indicate significant differences among WT, control, and *crr6* plants according to Tukey–Kramer multiple comparison tests (*p* < 0.05). black symbols represent WT and control plants, while orange symbols represent *crr6* mutant.

**Figure 5.**
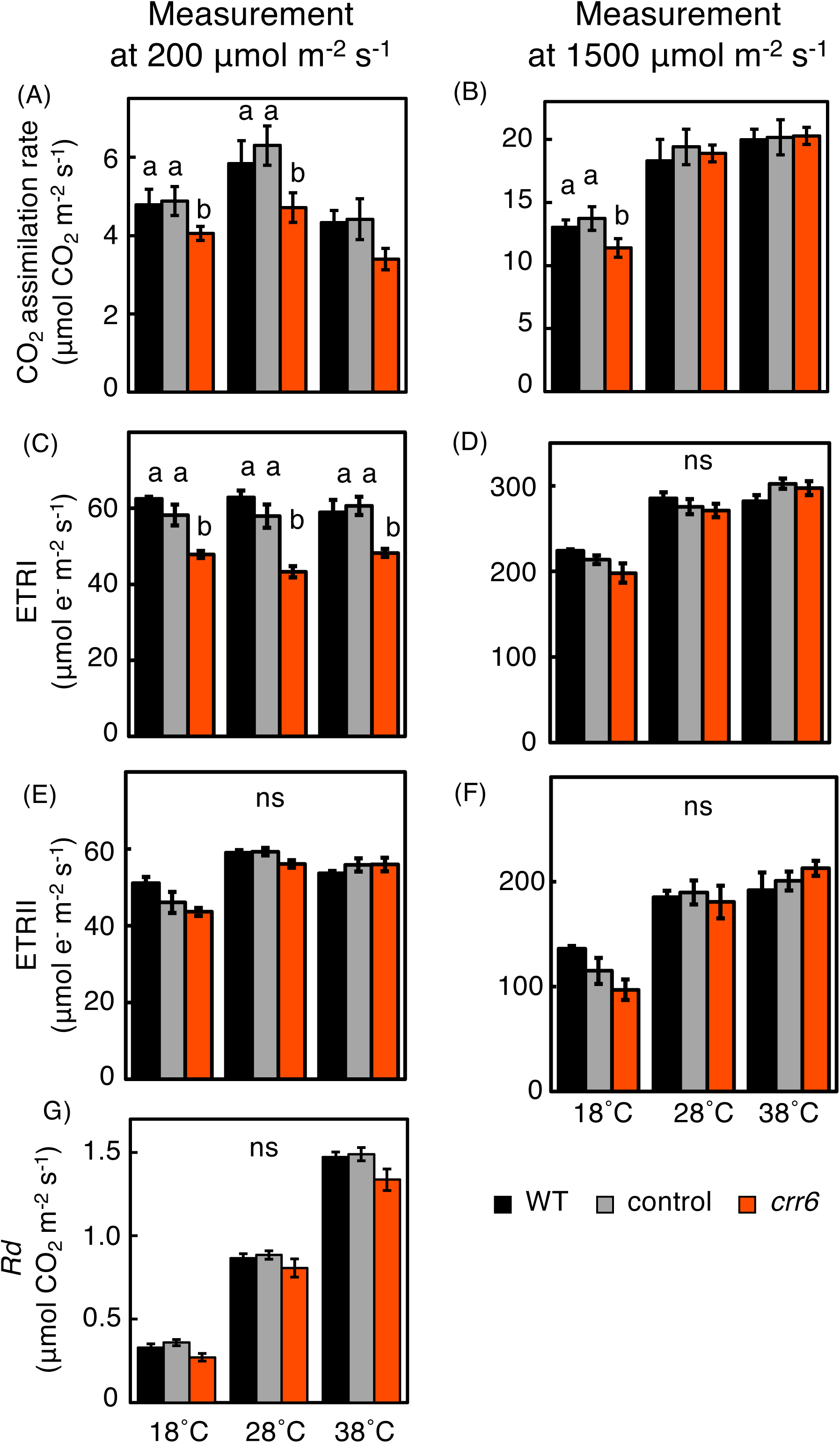
Comparison of photosynthetic parameters under low and high irradiance at different temperatures. Values of (A, B) net CO₂ assimilation rate, (C, D) electron transport rate through PSI [ETR(I)], (E, F) electron transport rate through PSII [ETR(II)], and (G) dark respiration rate (*Rd*) at each temperature were extracted from the light-response curves shown in Figs. 3 and 4 and compared between low light (200 μmol m⁻² s⁻¹) and high light (1500 μmol m⁻² s⁻¹). Values represent means ± SE (n = 3–5). Significant differences among genotypes were evaluated using Tukey–Kramer tests (*p* < 0.05). Different letters indicate significant differences among WT, control, and crr6 plants according to Tukey–Kramer multiple comparison tests (p < 0.05).

Consistent with this reduction in PSI electron transport, Y(NA) was elevated in the *crr6* mutant under sub-saturating light. Y(NA) increased under 750 µmol m⁻² s⁻¹ at 18 and 28°C and under 500 µmol m⁻² s⁻¹ at 38°C (Fig. 3G–I). Furthermore, the plastoquinone pool was more reduced in the mutant, as indicated by higher 1-qL values at 300–500 µmol m⁻² s⁻¹ at 18 and 28°C (Fig. 3M–O). NPQ and Y(ND) did not differ significantly between genotypes under these conditions (Fig. 3D–F, P–R). The ratio ETRI/ETRII, commonly used as an indicator of CET activity, was significantly lower in the crr6 mutant under low light at 28 and 38°C, suggesting reduced CET capacity.

Together, these results indicate that NDH deficiency leads to PSI acceptor-side limitation and enhanced reduction of the plastoquinone pool under relatively low irradiance, particularly at low temperature, thereby contributing to reduced CO₂ assimilation.

### NDH deficiency exacerbates photosynthetic decline under fluctuating light

Photosynthetic responses were examined under three fluctuating-light regimes (10-min HL/LL cycles, 5-min HL/LL cycles, 2-min HL/LL cycles) (Fig. 6). Initial photosynthetic parameters were consistent with the steady-state light-response measurements. Under fluctuating light, ETRI, ETRII, and CO₂ assimilation rates gradually declined in both genotypes; however, the decline was markedly greater in the *crr6* mutant during both high- and low-light phases (Fig. 6D, E, F, M, N, O, V, W, X). Consequently, final values of these parameters were substantially lower in the mutant than in WT plants.

**Figure 6.**
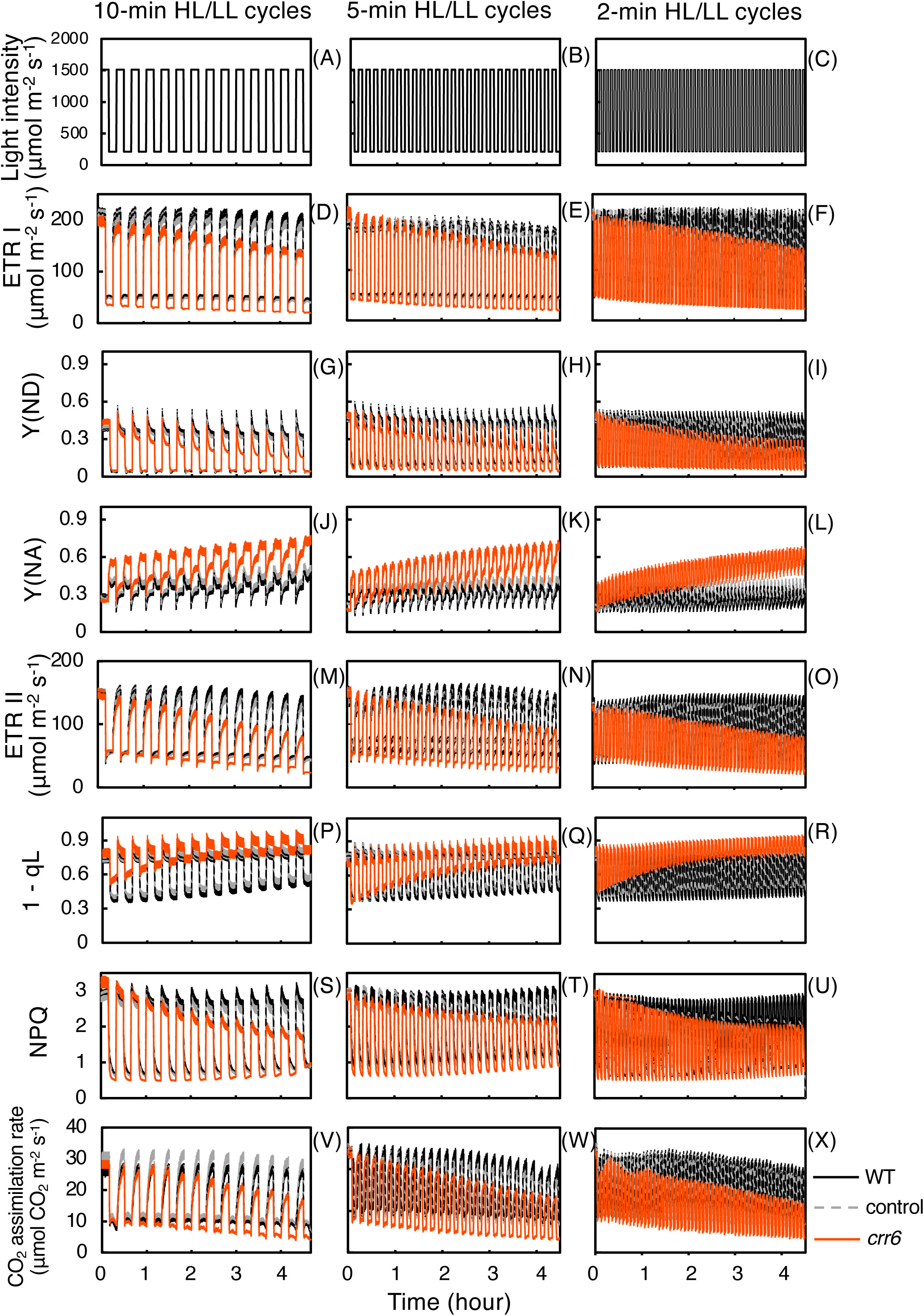
Photosynthetic responses to fluctuating light in WT and *crr6* plants. Photosynthetic parameters were measured at an ambient CO₂ concentration of 390 μmol mol⁻¹ under three fluctuating light regimes consisting of repeated cycles between high light (1500 μmol m⁻² s⁻¹) and low light (200 μmol m⁻² s⁻¹) for 4.5 h. Light fluctuations were applied at intervals of 10 min (A, D, G, J, M, P, S, V), 5 min (B, E, H, K, N, Q, T, W), or 2 min (C, F, I, L, O, R, U, X). Parameters shown are (A–C) measurement light conditions, (D–F) electron transport rate through PSI [ETRI], (G–I) donor-side limitation of PSI [Y(ND)], (J–L) acceptor-side limitation of PSI [Y(NA)], (M–O) electron transport rate through PSII [ETRII], (P–R) fraction of closed PSII centers (1 − qL), (S–U) non-photochemical quenching (NPQ), and (V–X) net CO₂ assimilation rate. Values represent means ± SE (n = 3–5).

PSI redox dynamics also differed between genotypes. In WT plants, Y(ND) and Y(NA) remained relatively stable throughout the treatment. In contrast, the *crr6* mutant showed a stepwise increase in Y(NA), reaching approximately 40% higher levels than WT under both light intensities (Fig.6J–L). Simultaneously, Y(ND) progressively decreased at 1500 µmol m⁻² s⁻¹ and ultimately fell to less than half of WT levels (Fig. 6G–I). Consistent with PSI over-reduction, the plastoquinone pool was more reduced in the mutant at both light intensities (Fig. 6P–R). NPQ induction was also suppressed, particularly during the high-light phases (Fig. 6S–U).

These findings indicate that NDH-dependent CET is essential for maintaining PSI redox balance and sustaining electron transport under fluctuating light, independent of fluctuation frequency.

### NDH-dependent CET protects PSI from photoinhibition under fluctuating light

To assess photoinhibition, maximum PSII quantum yield (Fv/Fm) and maximum P700 signal (Pm) were measured before and after 5 h of fluctuating light treatment (Fig. 7). Before treatment, both parameters were similar among genotypes. After fluctuating light exposure, Fv/Fm remained largely unchanged in WT and control plants. In contrast, Pm declined significantly in the *crr6* mutant, indicating selective PSI photoinhibition. Moreover, PSI photodamage became more severe as fluctuation frequency increased, demonstrating that NDH-dependent CET plays a critical role in protecting PSI under rapidly changing light environments.

**Figure 7.**
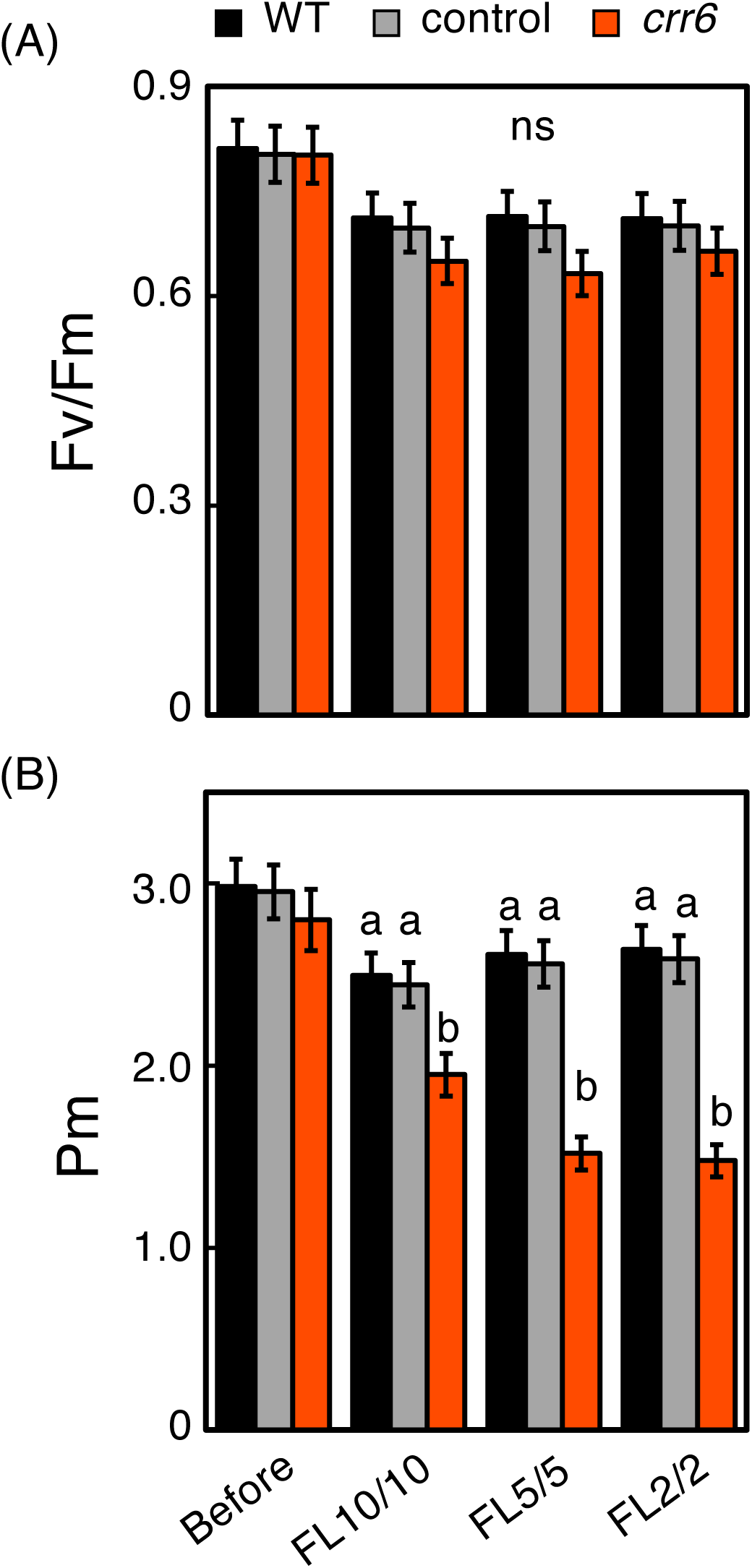
Photoinhibition of PSI and PSII following fluctuating light treatment in WT and *crr6* plants. Photoinhibition parameters were measured following the fluctuating light treatments shown in Fig. 6. Leaves were exposed to repeated cycles of high light (1500 μmol m⁻² s⁻¹) and low light (200 μmol m⁻² s⁻¹) for 4.5 h at intervals of 10, 5, or 2 min. After the treatment, leaves were dark-incubated for 30 min prior to measurements. (A) Maximum quantum yield of PSII (Fᵥ/Fₘ). (B) Maximum oxidizable P700 signal of PSI (Pₘ). Values represent means ± SE (n = 3–5). Different letters indicate significant differences among genotypes according to Tukey–Kramer multiple comparison tests (*p* < 0.05).

## Discussion

In natural field environments, plants are exposed to continuous fluctuations in irradiance and temperature that can constrain photosynthetic performance. However, the physiological significance of NDH-dependent cyclic electron transport under such realistic conditions has remained unclear. In this study, we addressed this question by analyzing NDH-deficient rice grown under field conditions. The mutant exhibited reduced CO₂ assimilation during fluctuating light treatments (Fig. 6), which was associated with significant decreases in biomass accumulation and grain yield (Fig. 2C–E). Physiological measurements further revealed that NDH deficiency impaired PSI electron transport and carbon fixation, particularly under sub-saturating irradiance and low temperature (Figs. 3, 4, 5). These findings indicate that NDH-dependent CET plays an important role in sustaining photosynthetic efficiency and plant productivity when crops experience dynamically changing environmental conditions in the field.

### NDH-dependent CET sustains PSI electron transport and carbon assimilation under weak light and low temperature conditions frequently encountered in the field

A central finding of this study is that NDH deficiency primarily impaired PSI electron transport and CO₂ assimilation under sub-saturating irradiance. At moderate (28°C) and high (38°C) measurement temperatures, the reduction in CO₂ assimilation in *crr6* mutant was confined mainly to the low-to-intermediate light range (Fig. 4A–C), whereas at low temperature (18°C) the impairment extended across nearly the entire light response curve. This temperature-dependent expansion of the phenotype strongly suggests that NDH-CET becomes increasingly important when biochemical carbon fixation capacity is restricted.

Importantly, this reduction in assimilation was not associated with changes in stomatal conductance, intercellular CO₂ concentration, respiration rate, or the abundance of major photosynthetic components such as Rubisco, chlorophyll, cytochrome *b*₆/*f*, or leaf nitrogen content (Fig. 4D–I; Fig. 5G; Table 1). Instead, the suppression of CO₂ assimilation closely paralleled the reduction in ETRI, indicating that NDH deficiency primarily limits the regeneration phase of the Calvin–Benson cycle through altered energy balance rather than diffusional or structural limitations. Because RuBP regeneration frequently limits photosynthesis under weak irradiance (Hikosaka et al., 2006), the loss of NDH-mediated *pmf* formation likely reduces ATP supply relative to NADPH, thereby constraining carbon fixation. This interpretation is consistent with previous reports demonstrating that NDH-dependent CET contributes to *pmf* formation by coupling ferredoxin oxidation to proton translocation across the thylakoid membrane (Strand et al., 2017) and supports photosynthesis under weak light in rice (Yamori et al., 2015). At low temperatures, the activity of Calvin–Benson cycle enzymes becomes kinetically restricted, which is one of the main factors promoting over-reduction of stromal electron acceptors and increasing susceptibility of PSI to photoinhibition (Sonoike, 2025). NDH-CET likely alleviates this imbalance by enhancing photosynthetic control at the cytochrome *b*₆/*f* complex and stabilizing PSI activity. Indeed, earlier work demonstrated that NDH-deficient rice shows decreased PSI activity and CO₂ assimilation under chilling conditions (Yamori et al., 2011), a phenomenon further supported by the broader light-range impairment observed here at 18°C.

Field environments are characterized by prolonged exposure to sub-saturating irradiance due to canopy shading, cloud movement, and diurnal solar geometry (Pearcy, 1990). Moreover, in temperate rice-growing regions, crops are frequently exposed to moderate or low temperatures throughout much of the growing season. Consequently, NDH-dependent CET is likely to play a particularly important ecological role under environmental scenarios where weak irradiance and low temperature co-occur. Because the duration of strong irradiance exceeding 1000 µmol m⁻² s⁻¹ is typically limited during a day, cumulative carbon gain in the field is largely determined by photosynthetic performance under sub-saturating light. The reduced biomass accumulation and grain yield observed in *crr6* plants (Fig. 2C–E) can therefore be interpreted as an integrated outcome of chronically reduced photosynthesis during these environmentally prevalent conditions. This interpretation is consistent with the well-established relationship between leaf photosynthetic capacity and plant productivity (Evans, 2013; Yamori et al. 2016b).

It should also be noted that alternative energy dissipation pathways may partially compensate for NDH deficiency. Previous studies suggested that enhanced photorespiration or activation of other cyclic or alternative electron transport routes can contribute to maintaining PSI redox balance when NDH function is impaired (Peterson et al., 2016; Chen et al., 2023). The relatively small difference in the CET activity index (ETRI/ETRII) between genotypes at low temperature observed here may support the possibility that additional regulatory mechanisms operate under such conditions. Further investigation will be required to clarify the detailed metabolic adjustments underlying the reduced assimilation in NDH-deficient plants.

### NDH-dependent CET stabilizes PSI redox balance and prevents cumulative photoinhibition under fluctuating light

Under simulated fluctuating light regimes, NDH-deficient plants exhibited progressive declines in ETRI, ETRII, and CO₂ assimilation (Fig. 6), accompanied by stepwise increases in acceptor-side limitation of PSI, as indicated by elevated Y(NA), and decreases in donor-side limitation Y(ND). These responses resulted in a selective reduction in the maximal oxidizable P700 signal (Pm) after fluctuating light treatment without a corresponding decrease in Fv/Fm (Fig. 7), clearly indicating PSI-specific photoinhibition.

Such vulnerability of PSI under fluctuating irradiance has been reported previously in NDH-deficient rice and Arabidopsis (Yamori et al., 2016a; Kono et al., 2017; Zhou et al., 2023). Mechanistically, CET pathways appear to play complementary roles during dynamic light transitions. The PGR5-dependent pathway rapidly induces ΔpH-dependent photosynthetic control under high light to restrict electron transport to PSI (Munekage et al., 2004; Suorsa et al., 2012), whereas NDH-dependent CET is likely to function as an electron sink receiving electron from Fd to prevent PSI acceptor-side excessively reduced (Takeuchi et al., 2025). From our results of light-response curve and other studies on the physiological role of NDH, this NDH function is more prominent during low-light phases when PSI acceptor-side limitations become critical due to delayed activation of stromal metabolism (Zhou et al. 2023; Yamori et al., 2015).

Interestingly, although the interval of light fluctuation did not markedly affect the magnitude of assimilation decline, PSI photoinhibition tended to become more severe as fluctuation frequency increased (Fig. 7). These findings suggest that cumulative carbon gain and PSI structural stability respond differently to temporal light dynamics. In natural field environments where irradiance can change on timescales of seconds, the ability of NDH-CET to facilitate rapid redox recovery of PSI acceptors may represent an important adaptive mechanism.

One important question is whether PSI photoinhibition accumulates in the natural environment. When we measured Pm before fluctuating light irradiation, Pm value was not different between WT and *crr6* mutant. PSI photoinhibition starts with a decrease of functional F_A_ and F_B_ clusters (Sonoike, 2025). However, this reduction did not affect P700 measurement (Tiwari et al., 2016). Therefore, it is possible that *crr6* mutant showed lower ETRI and CO₂ assimilation under a wide range of light intensity because of accumulated photoinhibition.

### Field environments integrate multiple environmental drivers that amplify NDH phenotypes

Natural terrestrial environments impose complex combinations of environmental constraints on photosynthesis (Yamori 2016). In addition to weak irradiance and temperature variation, spectral composition also changes dynamically. Light transmitted through vegetation or received during early morning and late afternoon is often enriched in far-red wavelengths, which preferentially excite PSI. Previous studies suggested that such spectral conditions can influence PSI excitation, PSI photoprotection, and NDH-dependent electron transport (Kono et al., 2017; Levin et al., 2026). Although spectral properties were not directly quantified in the present study, these findings imply that multiple environmental drivers may act simultaneously in the field to increase the physiological importance of NDH-CET.

Consistent with this ecological framework, NDH-deficient mutants exhibited substantial reductions in growth and yield under field cultivation despite showing relatively modest differences in biochemical leaf traits. These results indicate that the phenotypic impact of NDH deficiency likely reflects the cumulative effects of recurrent exposure to weak light, low temperature, fluctuating irradiance, and dynamic spectral environments. Under such realistic growth conditions, NDH-dependent cyclic electron transport appears to function as a key stabilizing mechanism that maintains long-term photosynthetic performance and crop productivity.

## Conclusion

This study demonstrates that NDH-dependent cyclic electron transport is required to sustain photosynthetic efficiency and productivity in rice grown under natural field conditions. NDH deficiency led to impaired PSI electron transport, reduced CO₂ assimilation, and consequent decreases in biomass and grain yield. These effects became particularly evident under environmentally relevant conditions such as weak irradiance, low temperature, and fluctuating light. Our findings indicate that NDH-CET contributes to maintaining photosynthetic robustness in dynamic terrestrial environments and represents an important regulatory component supporting crop performance under realistic growth conditions.

## Author contributions

H.K. and W.Y. conceived the study and designed the experiments. H.K. and W.Y performed the experiments and H.K. analyzed the data with contributions from W.Y. H.K. and W.Y. wrote the manuscript and approved the final version.

## Data availability

Supporting data can be requested by contacting the corresponding author.

## AI generative statement

Generative AI tools were used to assist with improving the clarity and language of the manuscript. All scientific content and interpretations were developed and verified by the authors. No AI tools were used to generate data, analyses, or figures.

